# You will know by its tail: a method for quantification of heterogeneity of bacterial populations using single cell MIC profiling

**DOI:** 10.1101/2022.04.29.490018

**Authors:** Natalia Pacocha, Marta Zapotoczna, Karol Makuch, Jakub Bogusławski, Piotr Garstecki

## Abstract

Severe non-healing infections are often caused by multiple pathogens or by genetic variants of the same pathogen exhibiting different levels of antibiotic resistance. For example, polymicrobial diabetic foot infections double the risk of amputation compared to monomicrobial infections. Although these infections lead to increased morbidity and mortality, standard antimicrobial susceptibility methods are designed for homogenous samples and are impaired in quantifying heteroresistance. Here, we propose a droplet-based label-free method for quantifying the antibiotic response of the entire population at the single-cell level. We used *Pseudomonas aeruginosa* and *Staphylococcus aureus* samples to confirm that the shape of the profile informs about the coexistence of diverse bacterial subpopulations, their sizes, and antibiotic heteroresistance. These profiles could therefore indicate the outcome of antibiotic treatment in terms of the size of remaining subpopulations. Moreover, we studied phenotypic variants of a *S. aureus* strain to confirm that the profile can be used to identify tolerant subpopulations, such as small colony variants, associated with increased risks for the development of persisting infections. Therefore, the profile is a versatile instrument for quantifying the size of each bacterial subpopulation within a specimen as well as their individual and joined heteroresistance.

## INTRODUCTION

Heterogeneity of an infecting bacterial population is an important consideration for the identification of an effective antibiotic treatment^1,2^. Some of the most common infectious diseases can be caused by more than one co-colonizing bacterial pathogen, incl. soft tissue infections, peritonitis, cystic fibrosis, urinary tract infections, and endocarditis^3^. An estimated 10% of antibiotic-resistant subpopulations is undetectable by current diagnostic tests leading to treatment failure^4^. Moreover, polymicrobial infections doubled the number of amputations in diabetic foot infections (DFI)^5^, while polymicrobial bloodstream infections were shown to increase the mortality rates from 24 to 47%^6^. Standard methods of antibiotic susceptibility testing (AST) fail to inform on the level of distribution of antibiotic resistance of the entire bacterial population of co-colonizing species. An example of co-infecting pathogens is *Staphylococcus aureus* and *Pseudomonas aeruginosa* abundant and prevailing in the airways and lungs of cystic fibrosis patients^7,8^. Inadequate antibiotic regimens are likely to interfere with the population’s complexity in an uncontrollable manner and may promote survival or emergence of resistant subpopulations or remove the non-pathogenic susceptible microflora^9^.

Moreover, in response to environmental stress bacteria can transiently modify their transcriptional profiles or acquire mutations. This type of stress-induced phenotypic heterogeneity has been known since the middle of the XX century^10^.These mutations often lead to the reduction in metabolic activity or changes to the cell membrane potential that result in an altered phenotype, such as a longer lag time, increased tolerance to antibiotics. Stress-induced phenotypic variants are also often associated with chronic infections^11–15^. Nicoloff et al reported that in ca. 27.4% of 766 bacteria-antibiotic combinations with 28 antibiotics, the bacterial strains were heteroresistant^16^. Moreover, heterogeneity of a bacterial strain in response to antibiotics has been shown to depend on changing the number of copies of antibiotic resistance genes. Gene copy number oscillation of plasmid- and chromosomally-encoded antibiotic resistance genes was shown responsible for changing the levels of resistance of bacterial cells within a bacterial population.^17,18^

Previous works have demonstrated the advantages of droplet microfluidics for the cultivation of bacteria and the determination of antibiotic breakpoints at the single-cell level^19–23^. Encapsulation of single bacterial cells with various antibiotic concentrations was used to determine the single-cell minimum inhibitory concentration (scMIC)^24^. However, due to heteroresistance, even a clonal population cannot be described by a single number, scMIC. Lyu et al.^25^ quantified bacterial heteroresistance by measuring the fraction of bacteria with MIC higher than antibiotic concentration c. In this paper, we denote the same fraction by F_R_(c) and refer to it as a ‘scMIC profile’. The scMIC profile is a function starting from 1 when all bacterial cells proliferate and monotonically decrease, reaching the value 0 when the antibiotic concentration is sufficiently high to inhibit the growth of all bacteria. The concentration corresponding to complete inhibition of the whole population determines ‘scMIC’, as defined by Artemova et al.^24^ The setup proposed by Lyu et al. for quantification of heteroresistance consisted of single bacterial cells encapsulated with antibiotic into ca. 75 pL droplets, which were then incubated under conditions optimal for bacterial growth following their probing with alamarBlue for detection of viability-dependent fluorescence signal in each droplet^25^. Lyu’s approach was a high-resolution method enabling detection of a resistant subpopulation as low as 10^−6^ of the entire population, yet the method required labeling, which limited its use. Similarly, a high-resolution droplet-based method for the determination of the scMIC and the distribution of scMIC was used by Scheler et al.^26^, who also went beyond a single cell level where the amount – not the concentration - of the antibiotic influences the bacteria inhibition. Heteroresistance profiles at a single cell level discussed by Scheler et al. referred to the probability distribution of MIC, p(MIC). The MIC distribution can be obtained from scMIC profile^26^ by taking the derivative, *p (c) =* -*dF*_*R*_(*c*)/ *dc*. Bacterial labeling was a factor that limited its experimental use to a single bacterial strain.

Recently, we reported that scattering can be used as a label-free detection of bacterial growth in nanoliter droplets and for the determination of scMIC^27^. Here, we extended the experimental setup by the implementation of an additional label-free detector of natural light emission which is specific to some bacterial species. We utilized the double detection device to investigate the scMIC profiles of phenotypically complex bacterial populations, incl. polymicrobial samples, and confirmed that the scMIC profiling efficiently informs antibiotic susceptibility and other parameters of population complexity, incl. the size of phenotypically diverse subpopulations. We hope that our model paves the way for a better characterization of antibiotic susceptibility in complex bacterial samples.

## RESULTS

### Single-cell MIC profiles of binary bacterial populations

We designed and built a microfluidic system for label-free detection of bacterial growth in nanoliter droplets to investigate scMIC profiles of polymicrobial samples. The optical setup shown in Supplementary Figure 1 was used for the simultaneous detection of scattered light as well as the native fluorescence within the 510-550 nm range. Bacterial strains differ in fluorescence excitation and emission spectra, and this feature can be used to distinguish them. The sensitivity of detection of autofluorescence was validated using droplets containing diverse bacterial species [Supplementary Figure 2 and Supplementary Table 1]. Native fluorescence was detected in all tested bacteria apart from isolates of *Staphylococcus spp*. Previously it has been shown that *S. aureus* produces autofluorescence in red spectral range^28^ (>600 nm, outside the detection bandwidth of our system) due to the presence of porphyrins^29^. At the same time, *e*.*g*., *Pseudomonas aeruginosa* emits a yellowish-green autofluorescence^28^ (within the detection bandwidth) from pyoverdine^30^.

We sought to use this label-free detection consisting of two selective detectors to determine the antibiotic susceptibility profiles of samples consisting of pre-mixed suspensions of *S. aureus* and *P. aeruginosa*. Both opportunistic pathogens are frequently co-isolated from infections of catheters, endotracheal tubes, skin, eyes, and the respiratory tract, incl. the airways of people with cystic fibrosis. A range of antibiotic concentrations was added to the pre-mixed suspensions prior to experiment. Emulsification of single bacterial cells from the sample into 1 nL droplets achieving ca. 10% of single inoculated droplets containing at least one cell capable of proliferation. In further discussion, we refer to these droplets as “non-empty”. Approx. 30 thousand droplets were produced from each sample. Readouts of scattered light and autofluorescence intensities were taken for each droplet after incubation for 18 hours at 37°C [Fig. 1a]. The number of droplets with multiplying bacteria was scored based on their high intensity of either signal and the fractions of positive droplets, f_+_, were calculated for each antibiotic concentration (see Supplementary Information for more details). We define scMIC by the concentration that inhibits all bacteria (here determined at the point where f_+_≤ 0.1%). The scMIC for the samples made of a pre-mixed population was determined at 1.5 µg/mL for ciprofloxacin and 4 µg/mL for tobramycin, which corresponded to the breakpoint concentrations of the less susceptible strain, as verified using the standard microdilution method [Fig. 1b]. It is worth mentioning that Artemova et al.^24^ determined scMIC in a similar way, although by plating cells on agar and counting colony growth. This could lead to different scMIC values due to the limited amount of antibiotic in droplets^26^. The application of an autofluorescence detector allowed for discrimination between bacterial subpopulations and for quantitative estimation of the proportion of either strain across antibiotic concentrations in a studied sample. Droplets containing *S. aureus* were detected by scattering whilst *P. aeruginosa* containing could be detected by both detectors. At the region of low antibiotic concentrations (up to 0.125 µg/mL for ciprofloxacin, or 0.5 µg/mL for tobramycin) the f_+_ values from scattered light corresponded to the growth of the entire mixed bacterial population [Fig. 1c-d]. The scMIC profiles determined by scattering and autofluorescence for ciprofloxacin were superimposed above the *S. aureus* breakpoint *c* of ca. 0.5 µg/mL, thus represented subpopulation of recoverable *P. aeruginosa* [Fig. 1c]. The scMIC profile for tobramycin above the 1 µg/mL was determined only from scattering as no autofluorescence could be detected above this concentration, suggesting the remaining recoverable cells were all *S. aureus*. We estimated the proportions of both species in the samples based on the fraction of positive droplets where bacteria were not treated with antibiotic. To obtain the sizes of *P. aeruginosa* populations we designated the ratios of the droplets recognized by high native fluorescence intensity to the positive droplets detected by the high signal of scattered light. We determined *P. aeruginosa* population at 50% ± 5% and 9% ± 1% for samples treated with ciprofloxacin and tobramycin, respectively. The remaining portions of the samples correspond to the sizes of *S. aureus* population.

**Fig. 1.**
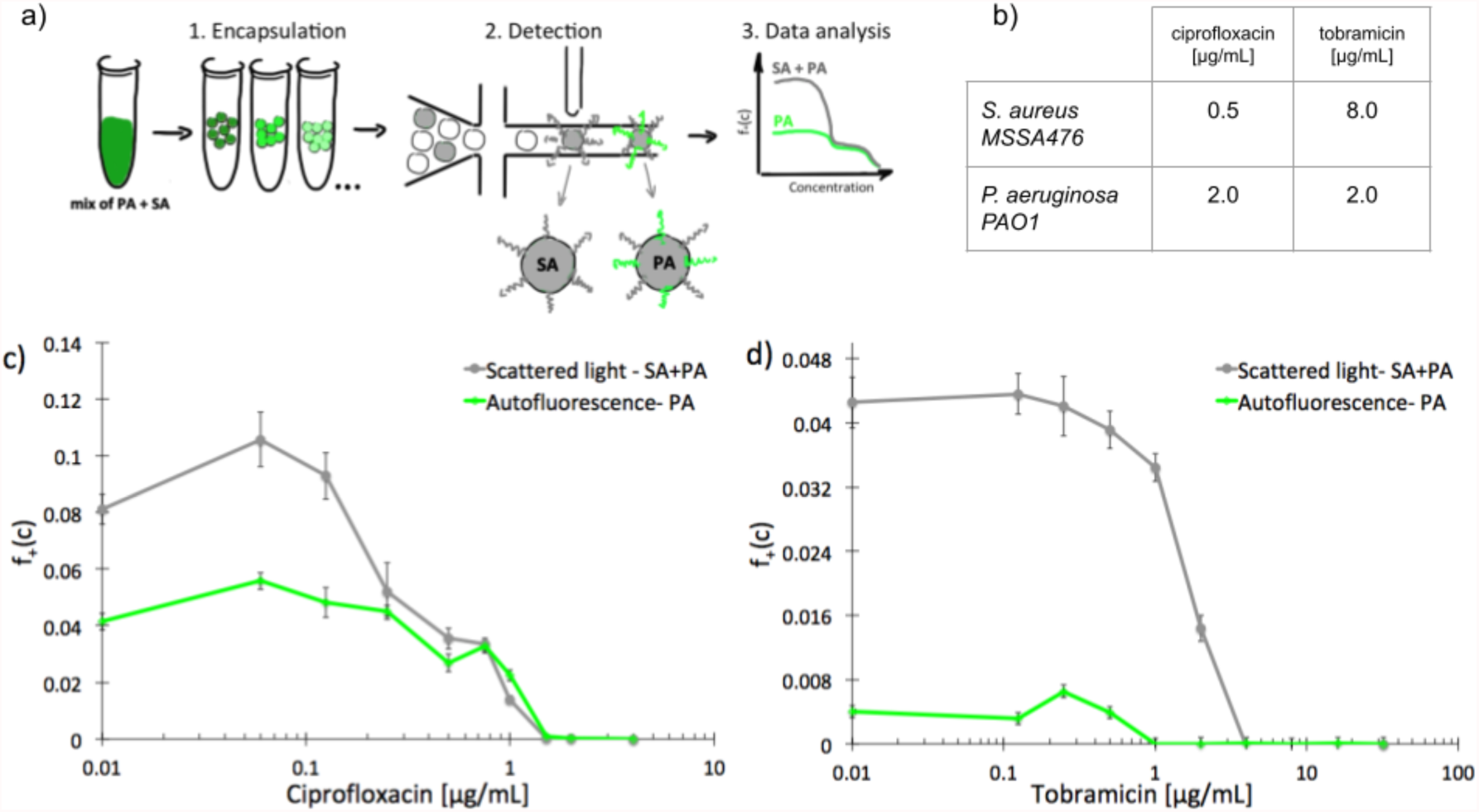
Experimental determination of scMIC profiles of polymicrobial samples of pre-mixed *S. aureus* MSSA47 and *P. aeruginosa* PAO1; a) schematic overview of the workflow for label-free determination of scMIC profile, b) MIC values determined by standard microdilution method for ciprofloxacin and tobramycin against studied strains, c-d) scMIC profiles of pre-mixed samples determined as a fraction of positive droplets, a function of antibiotic concentration measured based on scattered light (grey line) and autofluorescence (green line) intensity. Statistical treatment of error analysis is described in detail in Supplementary Information.

These results demonstrate the effectiveness of our method in the determination of the scMIC profile of a sample consisting of two bacterial subpopulations. Moreover, we observed that scMIC profiles obtained using scattering alone are sufficient to inform on the population complexity (existence of more than one subpopulation of bacteria with different level of antibiotic tolerance). Yet, in mixed populations of a large disproportion in subpopulation sizes (one strain over the other), autofluorescence is essential to discriminate.

### scMIC profiles of normal cells and small colony variants

Next, we used scattering-based detection to determine the scMIC profiles of phenotypically heterogeneous samples of a clonal strain. We selected normal morphology colonies (normal colony phenotype, NCP) of strain *S. aureus* SH1000 grown under optimal conditions, as well as colonies representing the small colony variant (SCV) phenotype, which we triggered with gentamicin [Fig. 2a]. Isolated SCV colonies were characterized by phenotypic changes, including aminoglycoside tolerance, small colony size, longer lag time, and reduced pigmentation [Fig. 2d, Supplementary Figure 3]. Both populations, consisting of cells harvested from either NCP or SCV colonies, were separately subjected to scMIC testing [Fig. 2b] and the respective scMIC profiles were determined (see Materials and Methods for more details). The NCP population required 2 µg/mL for the inhibition of the entire population while the scMIC of the SCV population was determined at 32 µg/mL [Fig. 2c]. Both scMIC values determined for either NCP or SCV were consistent with the breakpoint concentrations measured for the respective populations using the standard microdilution method [Supplementary Table 2].

**Fig. 2.**
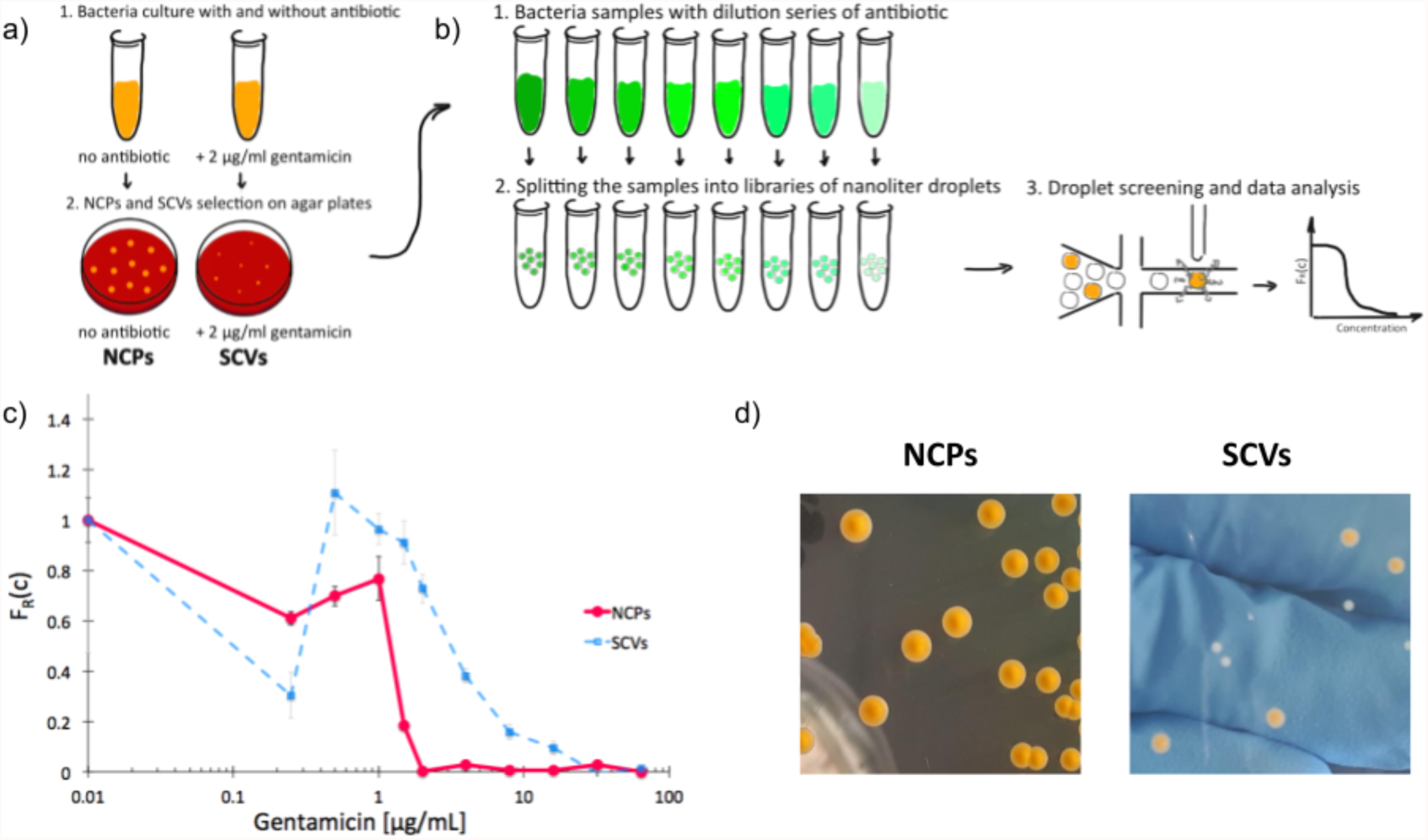
Experimental determination of single-cell MIC profile for *Staphylococcus aureus* of normal colony phenotype (NCPs) and small colony variants (SCVs); a) schematic overview of the experimental workflow of SCVs triggering and b) determination of scMIC incl. sample analysis towards phenotypic heterogeneity, c) comparison of single-cell MIC profiles determined for NCPs (red) and SCVs (blue), d) photographs of representative NCP and SCV colonies on tryptic soy agar, respectively. Statistical treatment of error analysis is described in detail in Supplementary Information.

Distinctive scMIC profiles, consisting of two regions, were observed for NCPs and SCVs [Fig. 2c]. The region of low antibiotic concentrations (up to gentamycin of ca. 1 µg/mL) with a slowly decaying fraction of recovering bacteria F_R_(c) was similar for both, NCP and SCV. The second region which we referred to as a transition region started at gentamicin of 1 µg/mL and was different in both populations. For NCPs the transition region ended at 2 µg/mL and had a sharp decline, while for SCVs it reached 32 µg/mL and had a much milder slope. Therefore, the transition region for SCVs was about 16 times broader than for NCPs. In this sense our SCVs population was 16 times more heterogeneous than NCPs, suggesting it consists of bacterial cells of larger distribution of antibiotic susceptibility. In the low antibiotic region with a high proportion of positive droplets the F_R_(c) values fluctuations were observed. They could be related to the decreased stability of non-empty droplets (droplets containing bacteria) likely to be caused by the cell aggregation, which is characteristic of the strain *S. aureus* SH1000.

### ‘Tails’ in the scMIC profiles

As we characterized the scMIC profiles of either NCPs or gentamicin-triggered SCVs, we wanted to investigate if they could determine their respective contribution within pre-mixed populations of a known proportion. The homogenous suspensions of NCPs and SCVs were combined into heterogeneous mixtures consisting of either 12% SCVs or 50% of SCVs and the remaining portion of NCPs, respectively [Fig. 3a]. As expected, the scMIC profiles of mixtures were combinations of profiles of either population. The profile of 12% SCV declined sharply in the transition region around 1 µg/mL to the F_R_(c) value of 0.08 and continued on with a minimal reduction in the shape of a tail between 1 and 8 µg/mL. The height of the tail at 0.08 directly corresponded to the size of SCV subpopulation and it amounts to 8% [Fig. 3b]. Whereas the scMIC distribution for the mixture of 50% SCV was broader, yet equally sharp starting at 2 µg/mL down to F_R_(c) ca. 0.37 and continuing as a tail ending at 16 µg/mL [Fig 3b]. Based on the height of the tail we estimated the SCV cells at 37% of the entire population. The difference between 50% and 37% follows from the precision of our method that we discuss in “Method precision” below.

**Fig. 3.**
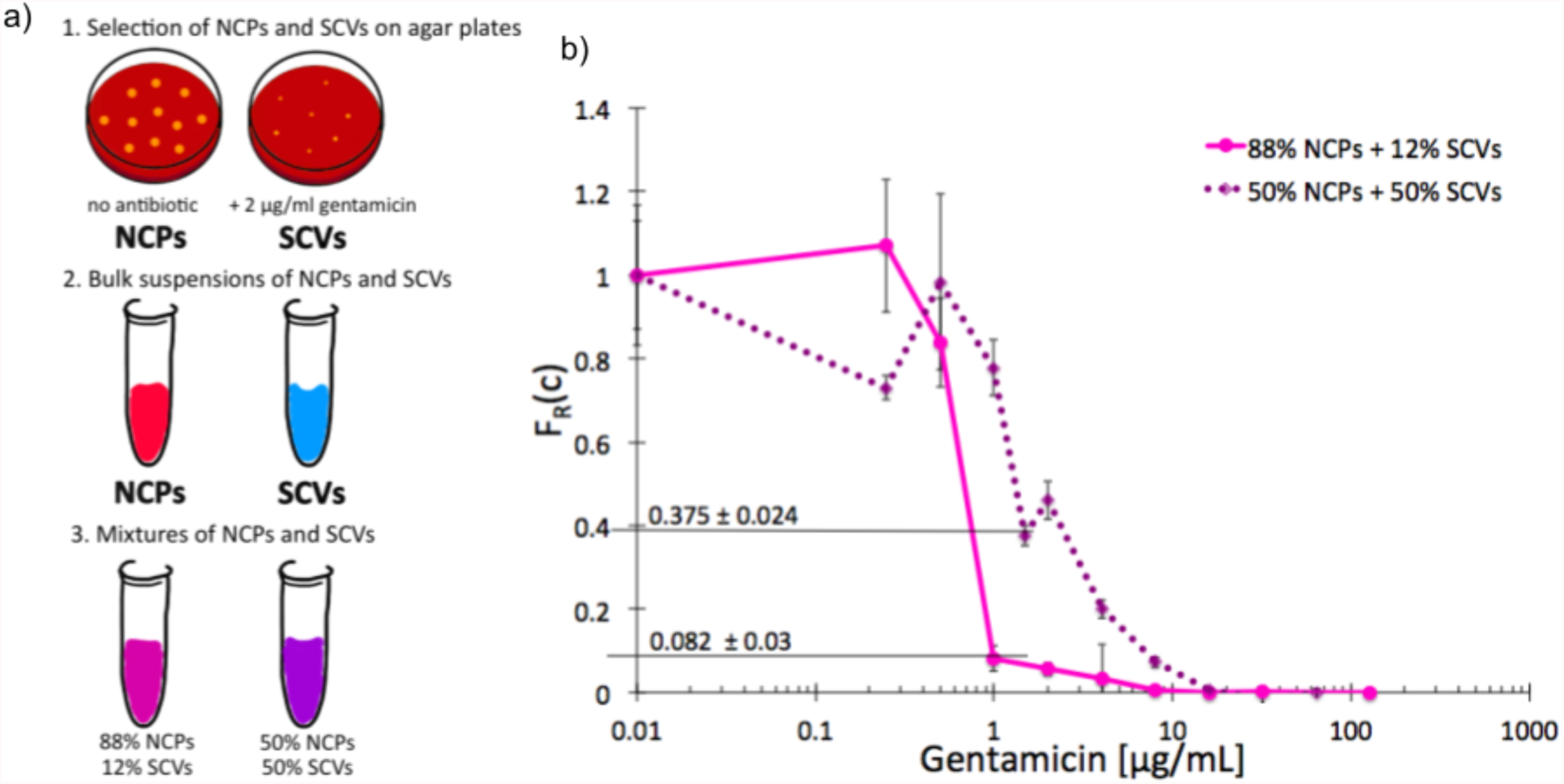
Single-cell MIC measurement of mixtures of phenotypic variants of *S. aureus* SH1000; a) schematic workflow of sample preparation, b) single-cell MIC profile of phenotypically heterogeneous mixtures containing respectively 12% (solid pink) and 50% (dotted purple) small colony variants (SCV) mixed with normal phenotype cells (NCPs). Statistical treatment of error analysis is described in detail in Supplementary Information.

These results indicate that the tail represents the size of SCVs subpopulation, whether it is low height corresponding to the low SCV content or high with a larger size subpopulation of tolerant bacteria. Consequently, the shape of scMIC profile is an indicator of coexistence of subpopulations of different distributions of antibiotic susceptibility and their respective sizes.

### “Tail” can inform phenotypic heterogeneity of a biofilm

The recalcitrance of *S. aureus* biofilms is commonly associated with the emergence of slow-growing subpopulations, such as the SCVs. We sought to investigate whether our label-free method is suitable for the detection of antibiotic tolerant subpopulations in a biofilm and to determine an adequate antibiotic dose for inhibition of this phenotypically heterogeneous population. We started by determining the unique scMIC profiles of NCPs and SCVs isolated from dispersed *in vitro* biofilm [Fig. 4a]. The scMIC of NCPs and SCVs were estimated at 1.5 and 32 ìg/mL, respectively, which corresponded to the scMIC profiles of either population isolated from suspension cultures [Fig. 2]. Moreover, the distinctive shapes of transition regions for either NCPs or SCVs were confirmed as a sharp decline and a milder slope, respectively [Fig. 4b]. Subsequently, we used scMIC profiles to estimate the size of SCV contribution in biofilms formed *in vitro*. The absence of a transition region (absence of a tail) in the scMIC profile of biofilm formed without gentamicin indicated it represents mostly a homogenous population. On the other hand, the biofilm formed in the presence of gentamicin had a transition tail with a base at 0.068 indicating the SCV content at 6.8% ± 1.9%, demonstrating the role of sub-MIC concentrations of gentamicin in triggering heteroresistance within a biofilm culture. [Fig. 4c,d].

**Fig. 4.**
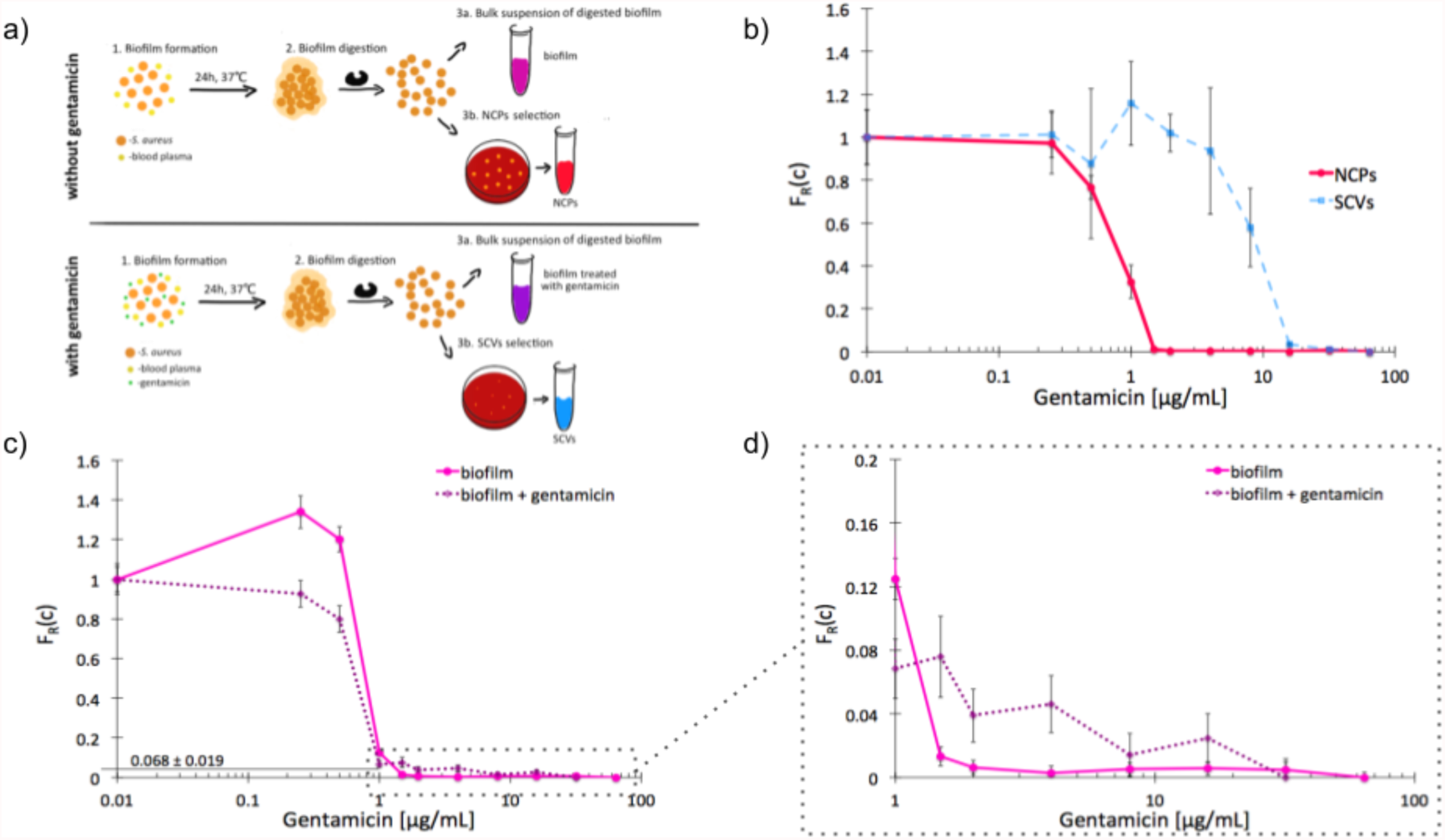
Single-cell MIC profiles of *S. aureus* dispersed from the biofilm. a) Schematic workflow of biofilms formation together with NCPs and SCVs selection in the absence (top) and presence (bottom) of gentamicin; b) heterogeneity profiles for NCPs (solid red) and SCVs (dotted blue) isolated from a biofilm cultured for 24 hours in the presence of gentamicin; c) scMIC profiles of biofilms cultured in the presence (dotted purple) and absence (solid pink) of gentamicin; d) the tails of scMIC profiles shown in c). Statistical treatment of error analysis is described in detail in Supplementary Information.

### Method precision

Measurements of scMIC profile require counting of droplets and determination of the number of positives. In an ideal experiment, we expect that the error of the fraction of resistant bacteria, 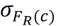, decreases as the square root of the number of non-empty droplets^26^, *Nf*_+_(0), as follows, 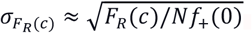. Therefore, one thousand non-empty droplets typically used in our experiments would give F_R_(c) fluctuations below 3%. The scMIC profile would be smooth - up to 3% fluctuation - monotonically decreasing curve. However, more significant fluctuations were observed in our experiments. For example, in Fig. 2c the fluctuations were sometimes larger than the error bars that we determined with the method described in Supplementary Information. These error bars have been determined under the assumption, that droplets in our experiments are stable, but we observed, especially with growing *S. aureus* SH1000 strain, that droplets have decreasing stability after incubation and tend to coalesce. For this reason, we estimated the accuracy of our method also by performing three repetitions of the same experiment. We cultured *S. aureus* biofilm for 72 hours in the presence of gentamicin. The biofilm was dispersed and subjected to scMIC experiments in three technical replicates measured one by one. Supplementary Figure 4 shows the heterogeneity profiles of all repetitions. Droplets without antibiotics or containing a low concentration of gentamicin were affected by more significant standard deviations. It is related to decreasing droplet stability with an increasing number of bacteria cells which we observed to be particularly visible for droplets containing *S. aureus* SH1000 cells as they tend to aggregate. These data show the highest deviation from the mean, 38%, in the transition region for concentration c=0.5 µg/ml. The remaining concentrations give the deviation of F_R_ smaller than 7%. Our numerical analysis revealed the following SCVs concentrations in repetitions no 1, 2, and 3: 17.9±1.4%, 31.3±2.9%, 29.8±3.0%, respectively. The average of obtained results is 26% and the standard deviation is 7.3%. Assuming a similar precision of the method (e.g. the relative error 7.3/26=28%) for about twice larger concentration (50% of SCV’s which corresponds to Fig. 3b) this precision explains the discrepancy between the prepared sample of 50% of SCV’s and the measurement based on a height of the tail (37%) in the discussion of Fig. 3b.

## DISCUSSION

Even in a clonal population, bacteria may differ from each other in their response to an antibiotic. On the one hand, a single bacterium in a droplet with an antibiotic can proliferate and colonize the droplet. On the other hand, the same antibiotic amount may inhibit the growth of another cell of the same bacterium. After incubation, the former situation leads to a droplet with plenty of bacteria (positive). In the latter case, the droplet contains a negligible number of bacteria (negative). F_R_(c) denotes the fraction of single-cell bacteria that “recover” from the treatment. In other words, F_R_(c) describes the inhibition of bacteria population at a single cell level which we refer to as the scMIC profile.

Here we proposed a dual label-free detection, which to the best of our knowledge, has never been reported before for quantification of bacterial growth in droplets. We found that the shape of the scMIC profile of a bacterial sample, estimated based on joint detection using scattering and autofluorescence, can inform about the coexistence of diverse subpopulations representing different autofluorescence (genus), sizes of respective subpopulations, the level of their respective antibiotic tolerance, and heteroresistance. These parameters are of key importance for the characterization of the response of complex bacterial populations to antibiotics. Although bacterial samples we investigated were derived from *in vitro* cultures, to achieve a controlled method validation, they were carefully selected to represent challenges in terms of diagnosis and treatment. We showed that the scMIC profile determined using our method was effective in sample discrimination between strains of *P. aeruginosa* and *S. aureus* in terms of their subpopulation sizes and antibiotic susceptibilities. Quantifying the size of subpopulations of co-infectious pathogens and their combined susceptibility to antibiotics, which may differ from individual ones, could be a breakthrough in infection control and make better use of existing antibiotic resources.

Similarly, we investigated scMIC profiles of small colony variants known to arise *in vivo* and promote chronic infection due to their altered phenotypes. Their transient nature presents a challenge for their identification and quantification, both in the clinics and research environment. Profiles of a population of SCVs triggered with gentamicin during exponential growth *in vitr*o demonstrated a large level of heterogeneity. This was reflected by the breath of the scMIC profile (16 times more heterogeneous response) of this population and suggests that the response of exponentially growing *S. aureus* to gentamicin which leads to the formation of SCVs may be in itself heterogeneous as it leads to different gene expression patterns resulting in antibiotic heteroresistance.

We found that populations containing SCVs could be identified by tails in their scMIC profiles. The shape of the scMIC profile tail, which we define as the region of scMIC profile starting at the bottom of a sharp decline of the fraction of recovering bacteria informed on the presence of SCVs. The tail height directly corresponded to the SCVs quantity thus confirming that the method was effective in the estimation of SCV subpopulations as small as 6,8%. To our knowledge, this is the first method that allows for quantitative estimation of this type of phenotypic variants which are of clinical relevance and associated with persistent infections.

Finally, the scMIC profile was informative in terms of the composition of a biofilm known for its heterogeneous nature, informing the size of either subpopulation within the biofilm culture. Application of our method could, therefore, inform on the absolute concentration of an antibiotic necessary to inhibit the growth of complex bacterial populations and provide an early stage indicator of the risk of heteroresistant pathogens associated with increased risks of complications.

## MATERIALS AND METHODS

### Bacterial strains and growth conditions

Bacterial strains of *Staphylococcus aureus* SH1000, *S. aureus* MSSA476 and *Pseudomonas aeruginosa* PAO1 were cultured in tryptic soy broth (Biocorp, Poland) for determination of scMIC. *Escherichia coli* ATCC25922, *Corynebacterium simulans* DSM 44392, *Staphylococcus intermedius* ATCC29663, *Staphylococcus aureus* Newman, *Staphylococcus epidermidis ATCC12228, Pseudomonas aeruginosa* ATCC27853, *Klebsiella pneumoniae* ATCC13883 and *Acinetobacter baumanii* ATCC19606 were cultured in lysogeny broth (Carl Roth, Germany) while *Salmonella arizonae* ATCC13314, *Shigella sonnei* ATCC29930 and *Listeria monocytogenes* ATCC19115 were cultured in brain heart infusion (Biocorp, Poland) for autofluorescence measurement.

### Triggering small colony variants

Small colony variants (SCVs) were triggered and isolated as described previously^31^. Briefly, tryptic soy broth containing gentamicin at 2 µg/mL was inoculated with a single colony of *S. aureus* SH1000 allowing culture growth at 37°C for 18h. SCVs were subsequently isolated by plating serial dilutions of the culture on tryptic soy agar plates not permitting the growth of normal cells but selecting for the spontaneous persistent variants of SCVs. The phenotypic switch was confirmed as smaller colony size, lack of pigmentation, and deficiency in beta-hemolysis. The SCVs colonies were harvested with a loop, suspended in 30% (v/v) glycerol solution, or directly subjected to experimental procedures described.

### *In vitro* biofilm formation

Biofilms were grown in microtiter plates (Nunclon™ Delta Surface, Thermo Scientific Nunc) either uncoated or pre-coated with human plasma. For plasma pre-coated wells a total of 100 µL of 20% (v/v) blood plasma in 50 mM carbonate buffer, pH 9.6, was incubated in the wells for 2h at 37°C prior inoculation with bacteria. Overnight cultures of *S. aureus* were diluted 1:200 in tryptic soy broth and inoculated at 100 µL in microtiter wells. Biofilms were formed for either 24h or 72h at 37°C. After washing off unattached cells, biofilm structures were dispersed with proteinase K, added to the wells at 100 µL for 30 min at 37°C. Digested biofilms were subjected to thorough pipetting following viable counts to determine their densities on either TSA or TSA containing 2 µg/mL of gentamicin. Gram-staining was performed to confirm that the aggregates were separated into single cells.

### Reagents

Chip fabrication: PDMS precursor and initiator (Dow Corning, USA) and tridecafluoro-1,1,2,2-tetrahydrooctyl-1-trichlorosilane (United Chemical Technologies, USA), Novec 1720 (3M, USA). Generation and reading of droplets: Novec HFE 7500 fluorocarbon oil (3M, USA) and triblock PFPE–PEG–PFPE surfactant. Antibiotics: gentamicin sulfate solution (TOKU-E, Japan and Sigma-Aldrich, USA), ciprofloxacin (TOKU-E, Japan), tobramycin sulfate (TOKU-E, Japan), vancomycin hydrochloride from Streptomyces orientalis (Sigma-Aldrich, USA), amikacin sulfate (Sigma-Aldrich, USA), rifampicin (TOKU-E, Japan), oxacillin sodium salt (Sigma-Aldrich, USA).

### Microfluidic Chip Fabrication

Microfluidic chips for droplet formation and detection were prepared in a four-step procedure. First, a polycarbonate (PC) mold is prepared by milling the microfluidic channels in 0.5 cm PC plate using a CNC machine (MSG4025, Ergwind, Poland). In the second step, polydimethylsiloxane (PDMS) is poured onto the PC chip, polymerized at 75 °C for 1 h, and the surface is activated by Laboratory Corona Treater (BD 20AC, Electro-Technic Products, USA) and silanized in the vapors of tridecafluoro-1,1,2,2-tetrahydrooctyl-1-trichlorosilane (United Chemical Technologies, USA) for 30 min under 10 mbar pressure. In the third step, the silanized mold is filled with liquid PDMS, polymerized at 75 °C for 1 h, and then the chips are bonded to 1mm thick glass slides using oxygen plasma. In the last step, the microfluidic channels are hydrophobically modified by introducing Novec 1720 (3M, USA), evaporation at room temperature, and baking the chips at 75 °C for 1 h.

### scMIC determination

The following assumptions were made to determine the scMIC profile, F_R_(c). For a given antibiotic concentration *c*, the same suspension of bacteria was encapsulated in *N*(*c*) droplets. They contained *N*_*ne*_(*c*) non-empty droplets (typically containing a single bacterium, the issue which we discuss below). After the incubation with an antibiotic of concentration *c*, they gave *N*_+_(*c*) positives. Thus, the fraction of recovering bacteria is given by *F*_*R*_(*c*) = *N*_+_(*c*)/ *N*_*ne*_(*c*). The number of droplets *N* and the number of positives *N*_+_ could be determined from the detector and they determine the fraction of positive droplets, *f*_+_(*c*) *= N*_+_(*c*)/*N*(*c*). But F_R_(c) required the number of non-empty droplets, *N*_*ne*_(*c*), as well. To determine the fraction of bacteria that recovered, F_R_(c), we used the experiment without antibiotic. Here, the number of non-empty droplets was equal to the number of positives, *N*_*ne*_(0) *= N*_*+*_(0), therefore, the fraction of non-empty droplets among all droplets is, *f*_*+*_(0) *= N*_*+*_(0)/*N*(0). Using the same bacteria suspension in the experiment with concentration c, we calculated the number of non-empty droplets by *N*_*ne*_ (*c*) = *f*_*+*_ (0) *N*(*c*). We then obtain, *F*_*R*_(*c*) = *f*_*+*_(*c*)/*f*_*+*_ (0). It is worth mentioning, that this relation between F_R_ and f_+_ does not take into account droplets with two, three, and more bacteria. However, as we explain in Supporting Information, in our experiments, most non-empty droplets contain a single bacterium, about 5% of them contain two, and 0.15% three bacteria. We also estimate, that neglecting droplets with two and more bacteria by using formula *F*_*R*_(*c*) = *f*_*+*_(*c*)/ *f*_*+*_(0), may change the value of *F*_*R*_ (*c*) by 10%.

The initial bacterial samples were diluted to the densities of 0.5 - 2·10^5^ CFU/mL providing a high probability of single-cell encapsulation (5-20% of positive droplets). Antibiotics were added to the sample at a range of concentrations the following emulsification into 1 nL droplets. The droplets were collected in 0.5 mL test tubes separately for each antibiotic concentration and incubated overnight at 37°C. After incubation, the droplets were scanned and the intensity of scattered or native fluorescence light was measured.

It was not always possible to determine from the detector signal (Supplementary Figure 2) whether the signal corresponds to a positive or to a negative droplet. It requires introducing a threshold that leads to a systematic error and an alternative approach to determine the scMIC profile. A detailed description of the determination of the scMIC profile with an explanation of all errors is shown in Figs. 1-4 (including the error related to the threshold) is in Supplementary Information.

### Scattered light intensity detection

Bacterial growth in droplets was measured based on scattered light intensity following the procedure described by Pacocha, Bogusławski *et al*. 2020. Positive and negative droplets were characterized by high and low intensity of scattered light, respectively. The number of positive compartments and a total number of droplets was used for the calculation of recovering bacteria fraction, F_R_(c), and scMIC profile determination.

### Autofluorescence intensity detection

Autofluorescence signal was detected in positive droplets by illumination with a 473 nm laser beam delivered by a fiber, collimated with a lens (focal length 19 mm), and focused by a 20× microscope objective. Native fluorescence light is collected by the objective, reflected on a dichroic mirror (a cut-off wavelength of 490 nm), passes a bandpass filter (central wavelength of 530 and 43 nm of bandwidth), pinhole, and targets a photomultiplier (Hamamatsu H5783-20). In this configuration, similar to a confocal microscope, only fluorescence originating from the focal volume will reach the photomultiplier. Thus, background autofluorescence (e.g., of PDMS) will be rejected. The optical setup shown in Fig. S1 was used for simultaneous detection of scattered and autofluorescence light intensity.

Negative droplets are visible due to the autofluorescence of culturing medium, while the presence of a large number of bacteria cells causes a pronounced increase in the fluorescence intensity. The intrinsic fluorescence of bacteria is linked with fluorescent amino acids (tryptophan, tyrosine, and phenylalanine) and coenzymes such as nicotinamide adenine dinucleotide hydride (NADH)^32^. It’s important to emphasize that the detection of bacteria from *S. aureus* genus is possible by optical setup modification. The excitation and spectrum of this bacteria lies in the UV spectral range (centered at approx. 280 nm and 330 nm, respectively)^32,33^. Thus, the detection of autofluorescence of *S. aureus* would require a different light source, and changing bandpass filter placed in front of the photomultiplier.

## ADDITIONAL INFORMATION

Supplementary Information accompanies this paper. The datasets generated during and/or analysed during the current study are available from the corresponding author on reasonable request.

## AUTHOR CONTRIBUTIONS

‡These authors contributed equally.

N.P. performed experiments, M.Z. designed the project, K.M. analyzed the data, N.P. and J.B. developed the autofluorescence-based detection of bacteria growth in the droplets, N.P., M.Z., K.M., J.B., P.G. wrote the manuscript.

## ACKNOWLEDGMENT

N.P. was supported by the National Science Centre funding based on decision numbers DEC-2014/12/W/NZ6/00454 (Symphony) and Maestro 10 2018/30/A/ST4/00036. M.Z. was funded by the National Science Centre, Poland grant nr 2018/31/D/NZ6/02648 (SONATA14), K. M. acknowledges NCN Maestro 10 2018/30/A/ST4/00036 and the support from the Foundation for Polish Science within the Team-Net POIR.04.04.00-0016ED/18-00 program, P. G. was supported by the Foundation for Polish Science (FNP) within TEAM TECH POIR.04.04.00-00-2159/16-00 and by The National Science Centre (NCN) within Maestro 10 2018/30/A/ST4/00036, J. B. was funded by the International Research Agendas program of the Foundation for Polish Science co-financed by the European Union under the European Regional Development Fund.

