## Supplementary Information for "You will know by its tail: a method for quantification of heterogeneity of bacterial populations using single cell MIC profiling"

**This PDF file includes:**

Figures S1 to S5
Tables S1 to S2
Supplementary text
SI References

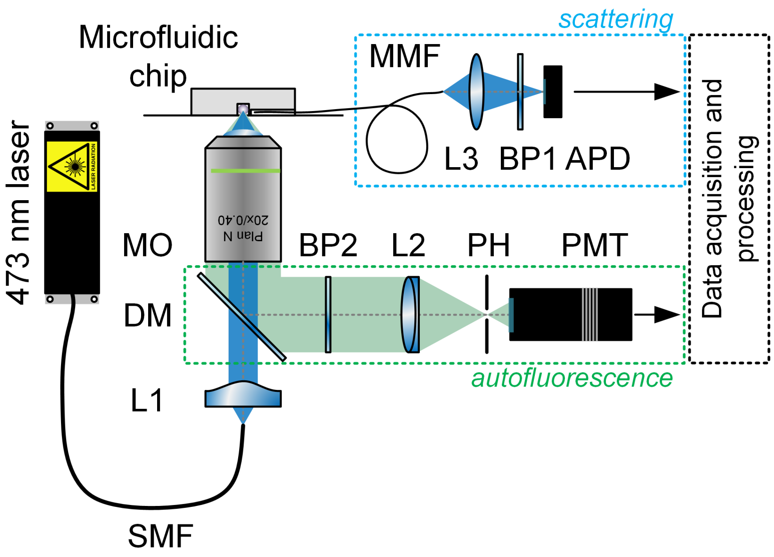


**Supplementary Figure 1.** Schematic representation of an optical setup for scattered light and native fluorescence intensity measurement. L1, L2, L3- lens, SMF – single-mode fiber, MMF- multimode fiber, BP1- bandpass filter (central wavelength of 470 and 10 nm of bandwidth) BP2- bandpass filter (central wavelength of 530 and 43 nm of bandwidth), DM- dichroic mirror (short-pass, cutoff wavelength of 490 nm), MO- microscope objective, PH- pinhole, APD- avalanche photodiode, PMT- photomultiplier.

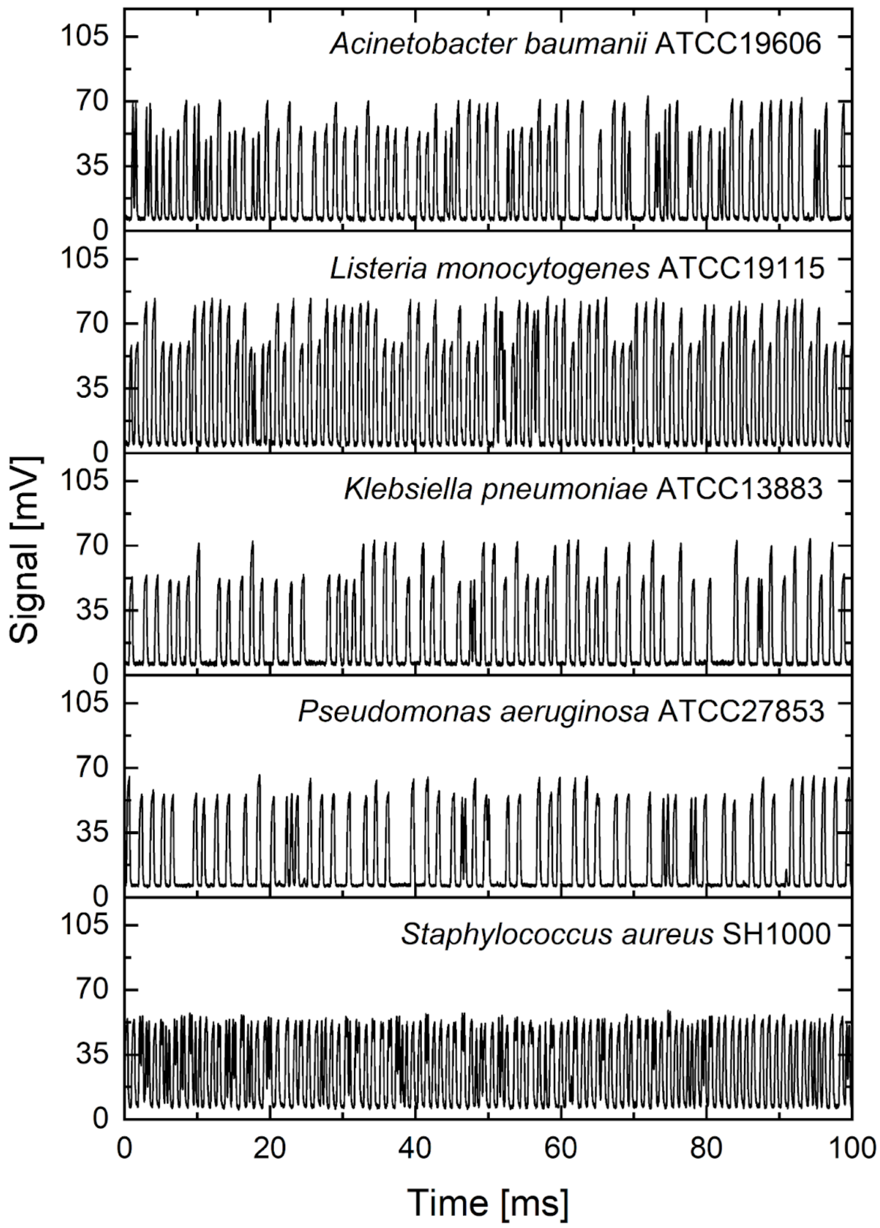


**Supplementary Figure 2.** Exemplary autofluorescence signal for samples containing 70% of negative droplets (growth medium) and 30% of positive droplets containing *Acinetobacter baumanii, Listeria monocytogenes, Klebsiella pneumoniae, Pseudomonas aeruginosa, Staphylococcus aureus* from top to bottom.

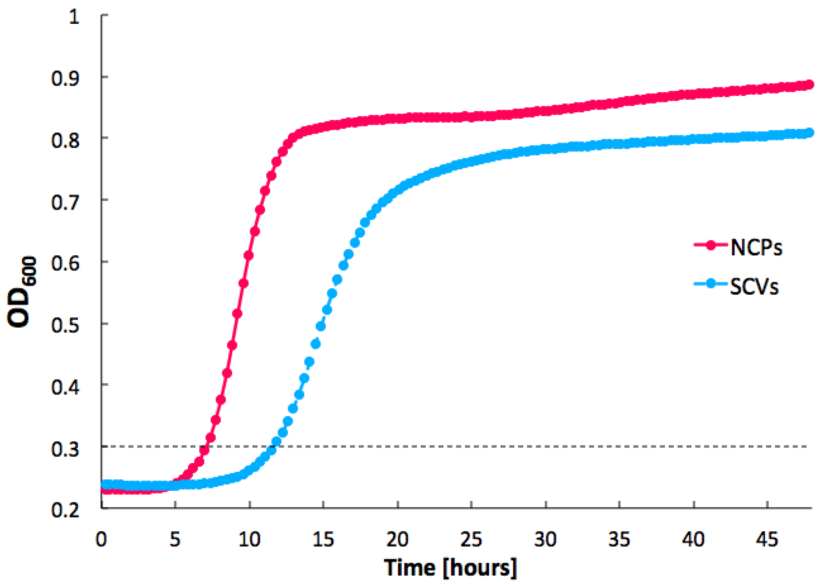


**Supplementary Figure 3.** Growth curves of normal colony (pink line) and small colony variant (blue line) phenotypes cultured in rich medium measured at 600 nm wavelength over 46 hours.

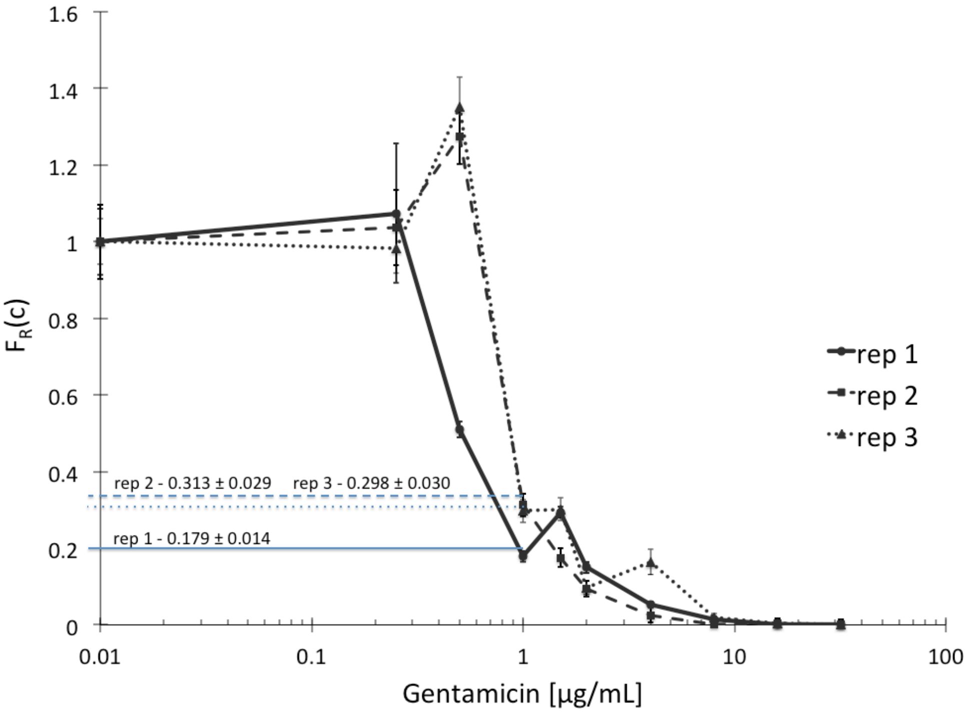


**Supplementary Figure 4.** Precision of scMIC curve determination. The graph presents heterogeneity profiles of dispersed *S. aureus* biofilm of 72h cultured in tryptic soy broth with gentamicin, analyzed in three repetitions. X and Y-axis indicate concentration of gentamicin and fraction of recovering bacteria, respectively.

**Supplementary Table 1.** Bacterial strains tested towards native fluorescence intensity allowing their label-free detection in nanodroplets. The ratio of positive to negative droplet intensities indicates the capability of droplet recognition and informs quantitative difference between both droplet populations. The dash stands for even intensities of positive and negative compartments.

| Bacterial species | Positive to negative peak ratio |
|---|---|
| *Escherichia coli* | 1.304 ± 0.109 |
| *Klebsiella pneumoniae* | 1.368 ± 0.007 |
| *Acinetobacter baumanii* | 1.294 ± 0.015 |
| *Pseudomonas aeruginosa* | 1.176 ± 0.005 |
| *Salmonella arizonae* | 1.116 ± 0.005 |
| *Listeria monocytogenes* | 1.357 ± 0.010 |
| *Shigella sonnei* | 1.0913 ± 0.0004 |
| *Enterococcus faecalis* | - |
| *Staphylococcus aureus Newman* | - |
| *Staphylococcus aureus SH1000* | - |
| *Staphylococcus epidermidis* | - |
| *Staphylococcus intermedius* | - |

**Supplementary Table 2.** Breakpoint concentrations of normal colony phenotype (NCP), small colony variant (SCV) and half on half mixture of both phenotypes to antibiotics of various types using standard microdilution method.

| | **Gentamicin** [µg/mL] | **Vancomycin** [µg/mL] | **Amikacin** [µg/mL] | **Rifampicin** [µg/mL] | **Oxacillin** [µg/mL] |
|---|---|---|---|---|---|
| NCPs | 2 | 2 | 16 | 0.0156 | 0.25 |
| 50% NCPs + 50% SCVs | 32 | 2 | 256 | 0.0156 | 0.25 |
| SCVs | 32 | 2 | 256 | 0.0156 | 0.25 |

**Supplementary Information Text**

**Determination of single cell minimum inhibitory concentration profiles.** Supplementary Figure 2 shows examples of signals from the detector. The first stage of the analysis is to identify droplets in the signal. For each signal corresponding to thousands of droplets, we manually and randomly pick up part of the signal that corresponds to a droplet that seems empty. We denote the part of the signal corresponding to the chosen droplet by $s_{empty}(t)$, which is defined for $t \in [0, T_d]$. The value of $T_d$ is typically equal to $0.0003s$. It is a duration of a passage of a droplet through the detector window. To facilitate the analysis we shift it by $s_{d0}$ and normalize it, $D(t) = \left(s_{empty}(t) - s_{d0}\right)/\eta$, such that $\int_0^{T_d} d\,t'\,D(t') = 0$ and $\frac{1}{T_d}\int_0^{T_d} d\,t'\,D(t')D(t') = 1$. We then calculate the convolution of the signal coming from the chosen droplet with the whole signal as follows,

$$C(t) = \frac{1}{T_d}\int_0^{T_d} d\,t'\,s(t + t')D(t').$$

From the above definition of the convolution and from described properties of $D(t)$, it follows that the convolution $C(t)$ is approximately equal to 0 for a constant signal. Moreover, the convolution $C(t)$ will have maxima for $t$ corresponding to droplets' beginnings. By the identification of maxima in $C(t)$ we identify droplets in the signal. Their number is denoted by $N_d$.

To verify whether this procedure leads to proper identification of droplets, we randomly chose about 56 droplets among about 10 thousand in each experiment. More specifically, we chose 8 sets of the signal, each with 7 identified droplets by the computer python custom code. We performed this procedure for every experimental point shown in the article (around 120 signals with about ten thousand droplets each). We noticed that it sometimes happens that droplets are not correctly identified (e.g. the procedure ignored a droplet that gives much smaller signal in comparison to a typical droplet). It introduces an error in the determination of the number of droplets.

We determine the error of the number of droplets, $\sigma_{N_d}$, by randomly choosing a set of $n_{df} = 7$ subsequent identified droplets. We count the number of droplets manually in each of $N_F = 8$ sets. The manually counted numer of droplets is denoted by $n_{realDF}$. We quantify the difference between the number of automatically and manually detected droplets by the ratio

$$r_F \equiv \frac{n_{realDF} - 1}{n_{df} - 1}$$

defined for each set. Dispersion of $r_F$ is denoted by $\sigma_{\langle r_F \rangle}$. The above-defined $r_F$ is interpreted as the ratio of manually identified droplets to the number of automatically identified droplets by the code,

$$N_{real} = N_d \langle r_F \rangle.$$

From the error propagation formula, we calculate the standard deviation of $N_{real}$, obtaining $\sigma_{N_{real}} = \sigma_{\langle r_F \rangle} N_d$. Using the fact, that $\langle r_F \rangle$ is typically close to unity, this formula also determines the error of the number of droplets

$$\sigma_{N_d} = \sigma_{\langle r_F \rangle} N_d.$$

When in all $N_F$ measurements we obtain $r_F = 1$, we estimate $\sigma_{\langle r_F \rangle}$ by the error in the situation in which our code makes a mistake in one droplet only. For $N_F\left(n_{df} - 1\right)$ droplets, the estimation for the error is (Poisson process [1])

$$\sigma_{\langle r_F \rangle} = \frac{1}{\sqrt{N_F\left(n_{df} - 1\right)}}.$$

Next, we calculate intensities of each identified droplet by

$$I = \frac{1}{T_d} \int_0^{T_d} d\,t\,(s(t_{ce} + t) - s_0),$$

where $s_0$ is the background signal level (the signal level outside the droplet) and $t_{ce}$ is the time corresponding to the beginning of a droplet in signal $s(t)$. We make histogram of the intensities. We typically observe two types of histograms. The first type of histogram has a gap that separates empty and positive droplets (e.g. the histogram correspoding to $0\mu g/ml$ in left panel of Supplementary Figure 5. In the second type of histogram, the intensities coming from positive droplets form a tail to the right of empty droplets (the histogram corresponding to $2\mu g/ml$ in the right panel of Supplementary Figure 5). In the former case, it is clear that the threshold that separates positive from negative droplets should be put in the gap. Here, the threshold position does not change the number of positives as long as it is in the gap. The situation is more complicated in the latter case because changing the threshold varies the number of positive droplets. Here, the distribution of intensity of positives partially overlaps with the distribution of negatives. However, under not too restrictive assumptions, it is possible to determine scMIC profile in both cases within one procedure. To explain the procedure, let's focus on the case with a tail.

To analyze histograms, we introduce the fraction of droplets with higher intensities than $I$ and denote it by $\alpha(I,c)$. It is thus defined by

$$\alpha(I,c) \equiv N_{aboveI}(I,c)/N_d(c),$$

where $N_{aboveI}(I,c)$ is the number of identified droplets with higher intensities than $I$ in the experiment with antibiotic concentration $c$. Because, $N_{aboveI}(I,c)$, is the result of counting, its error is determined as in the Poisson distribution [1] and is given by $\sigma_{N_{aboveI}} = \sqrt{N_{aboveI}}$. Using the error propagation formula in the above equation we get

$$\sigma_\alpha = \left[\frac{\alpha(I,c)}{N_d(c)} + \left(\alpha(I,c)\frac{\sigma_{N_d}}{N_d(c)}\right)^2\right]^{1/2}.$$

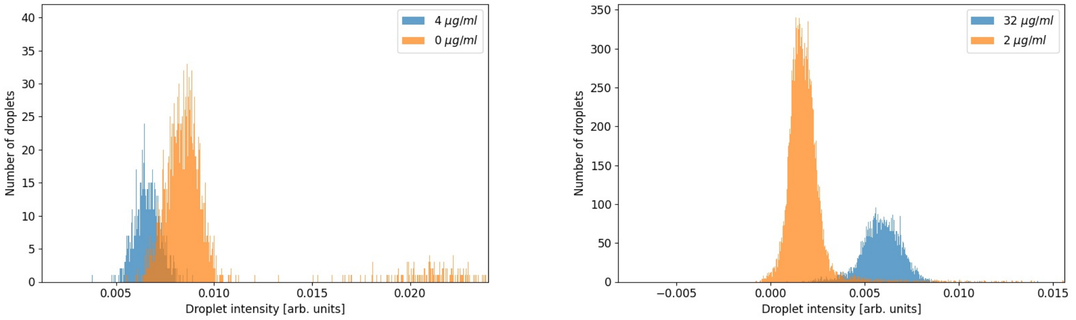


**Supplementary Figure 5.** Example of histograms of droplet intensities for two different antibiotic concentrations in a single experiment (left panel). In the right panel histograms correspond to another experiment.

Histograms contain intensities corresponding to positive and negative droplets, so they are composed of two distributions. We denote the probability density of intensities of empty droplets by, $p_{int}^{-}(I,c)$, while for positves by, $p_{int}^{+}(I,c)$. We, therefore, expect that the intensity distribution is given by a mix of positive and negative droplets,

$$p(I,c) \quad = \left(1 - xF_R(c)\right)p_{int}^{-}(I,c) + xF_R(c)p_{int}^{+}(I,c).$$

Here, $x$ is the fraction of nonempty droplets and $F_R(c)$ is the fraction of resistant bacteria in the population. Integrating the above equation from $I_{lim}$ to infinity we get,

$$\int_{I_{lim}}^{\infty} d\,I\,p(I,c) = \left(1 - xF_R(c)\right)\int_{I_{lim}}^{\infty} d\,I\,p_{int}^{-}(I,c) + xF_R(c)\int_{I_{lim}}^{\infty} d\,I\,p_{int}^{+}(I,c). \quad (1)$$

The above equation allow us to calculate the ratio $F_R(c)/F_R(0)$ and using the fact that $F_R(0) = 1$ we obtain,

$$F_R(c) = \frac{\alpha(I_{lim},c) - \alpha_-(I_{lim},c)}{\alpha(I_{lim}^0,0) - \alpha_-(I_{lim}^0,0)} \times \frac{\alpha_+(I_{lim}^0,0) - \alpha_-(I_{lim}^0,0)}{\alpha_+(I_{lim},c) - \alpha_-(I_{lim},c)}, \quad (2)$$

$$\alpha(I_{lim},c) \equiv \int_{I_{lim}}^{\infty} d\,I\,p(I,c),$$
$$\alpha_+(I_{lim},c) \equiv \int_{I_{lim}}^{\infty} d\,I\,p_{int}^{+}(I,c),$$
$$\alpha_-(I_{lim},c) \equiv \int_{I_{lim}}^{\infty} d\,I\,p_{int}^{-}(I,c).$$

In the above, we used $I_{lim}^0$ for Eq. (1) for the case $c = 0$. If we can measure $\alpha(I,c)$ and $\alpha_{-/+}(I,c)$, then the above equation would be sufficient to determine $F_R(c)$. However, we only showed how to measure $\alpha(I,c)$ and its error. Let's notice that the second factor in (2) should not depend on concentration $c$ when the distribution density of positives and negatives do not change with the concentration. We expect these conditions in our experiments. As a consequence and by choosing

$$I_{lim} = I_{lim}^0$$

we eliminate the second factor in Eq. (2) obtaining

$$F_R(c) = \frac{\alpha(c) - \alpha_-(c)}{\alpha(0) - \alpha_-(0)}. \quad (3)$$

To use this formula, we need to measure the distribution of negatives, $\alpha_-(I,c)$. We measure it in the experiment with the highest antibiotic concentration, which we always take to be sufficiently high to inhibit the growth of any bacteria.

The above point requires a comment. Let's discuss the histograms in the right panel of Supplementary Figure 5. If we performed the measurements of droplet intensities in the same conditions (e.g., exact droplet sizes, same laser intensity, droplet stability), we would expect that the maxima of both histograms overlap. But it is not the case: we observe a shift and a rescaling of signals. The shift and rescaling are consistent with the observation that the background levels of corresponding detector signals differ by a factor 4/3. The observed shift and rescaling violate our assumptions that $\alpha_{-/+}(I_{lim},c)$ does not depend on concentration $c$. Therefore, the second factor in (2) does not drop out. But it is possible to choose different $I_{lim}$ for the concentration $c$ to eliminate this factor. We observe that the signal coming from the negative droplets for concentration $c$ (typically Gaussian-like shape) is rescaled with respect to the highest concentration $c_{HAC}$ as follows,

$$p_{int}^{-}(I,c) = \frac{1}{r}p_{int}^{-}(rI - I_0, c_{HAC}).$$

The parameters $r$ and $I_0$ are determined for antibiotic concentration $c$ by the following fitting procedure. We assume that the positive droplets do not contribute to the lowest intensities. It allows us to use droplets with the lowest intensities (typically 80% of droplets) to make the fit of $p_{int}^{-}(rI - I_0, c_{HAC})/r$ to the distribution $p(I,c)$ obtained from the histogram. By the above fitting procedure one determines $I_{lim} = r(c)I_{lim}^0 - I_0(c)$ which eliminates the second factor in Eq. (2) obtaining again expression (3).

We represent this expression in the following form

$$F_R(c) = \frac{M(c)}{M(0)},$$

where

$$M(c) \equiv \alpha(c) - \alpha_-(c),$$

which we use to determine scMIC profiles, $F_R(c)$, in our article. We determine the error of $F_R(c)$ from the error propagation formula as follows,

$$\sigma_{F_R}(c) = \sqrt{\left(F_R(c)\frac{\sigma_M(c)}{M(c)}\right)^2 + \left(F_R(0)\frac{\sigma_M(0)}{M(0)}\right)^2},$$

where

$$\sigma_M(c) = \sqrt{\left(\sigma_{\alpha(c)}\right)^2 + \left(\sigma_{\alpha_-(c)}\right)^2}.$$

The errors shown in the paper are determined with the above procedure unless explicitly stated. The above method depends on $I_{lim}$, which introduces a systematic error. However, in all cases, we modified $I_{lim}$ to check whether it influences the results. We obseved that the changes of $F_R(c)$ with variation of $I_{lim}$ were within errors of scMIC, $\sigma_{F_R}(c)$. Therefore, we conclude that the systematic error due to threshold $I_{lim}$ does not influence scMIC profiles shown in the paper.

**Error coming from assumption that in each non-empty droplet there is only one bacterium.**

Suppose that in experiments we have only empty droplets and droplets containing a single bacterium. It is then straightforward to determine the fraction of resistant bacteria. Let $N(c)$ denotes the number of droplets in the experiment with concentration $c$ and $N_+(c)$ is the number of positives. Droplets in experiments with different antibiotic concentrations are emulsified from the same bacteria suspension. Therefore, in all experiments the fraction of droplets that contain a bacterium is the same and can be determined from the experiment without antibiotic, $f_+(0) = N_+(0)/N(0)$. The fraction of resistant bacteria, $F_R(c)$, by definition equals to the number of bacteria that can proliferate (leading to positive droplets), $N_+(c)$, divided by the number of all bacteria in the population (droplets that contain a single bacterium), $f_+(0)N(c)$. Thus $F_R(c) = f_+(c)/f_+(0)$, where $f_+(c) = N_+(c)/N(c)$ is the fraction of positives in the experiment with antibiotic concentraction $c$.

Here we estimate the error of our measurement that comes from the fact that we neglect droplets with two or more bacteria. For the estimation, we make two assumptions. We assume that in the process of emulsification of bacteria, they are randomly closed in droplets. Moreover we assume that two and more bacteria in droplets do not influence their proliferation. These situation have already been discussed by Scheler *et al.* [2] and leads to the following expression for the fraction of resistant bacteria, $F_R(c) = \log(1 - f_+(c))/\log(1 - f_+(0))$. By performing Taylor expansion of the logarithms we observe, that the above expression becomes $F_R(c) \approx f_+(c)/f_+(0)$ in the limit of small $f_+$. Moreover, the correction to the formula $F_R(c) \approx f_+(c)/f_+(0)$ is of the order of $F_R(c)f_+(0)$. Because $f_+(c)$ is typically 10% in our experiments, we neglect this correction in our analysis.
